# Ontogeny of vocalizations in Adlie penguin (*Pygoscelis adeliae*) chicks

**DOI:** 10.64898/2026.01.23.701401

**Authors:** Michele L. Adams, Danielle T. Fradet, Megan A. Cimino, Easton R. White, Laura N. Kloepper

## Abstract

Passive acoustic monitoring (PAM) is an efficient method to monitor dense aggregations of vocal animals but requires understanding the acoustic ecology of the species under examination. Avian vocal development is largely understood from songbirds, limiting its application to non-passerine taxa with different social and environmental pressures. As an example, colonial seabirds such as the Adélie penguin (*Pygoscelis adeliae*) inhabit acoustically crowded environments and rely on vocal cues, in addition to spatial information, for parent-offspring recognition. While adult penguin vocal communication is well studied, chick vocal development remains poorly characterized. Using the deep learning-based system DeepSqueak, we aimed to characterize the vocal development of wild *P. adeliae* chicks in the West Antarctic Peninsula. We found that acoustic features of chick calls changed systematically with age, with calls becoming longer and more frequency modulated over time. Characterizing chick vocal development from hatch to fledge provides important information to study phenological communication patterns in vocal-dependent seabirds and supports the application of PAM to assess climate-driven impacts on indicator species.

## Introduction

Vocal learning pertains to the capacity of specific mammal and bird species to learn and adjust their vocalizations through imitation and social interaction (Brenowitz & Beecher, 2005; Elie & Theunissen, 2020). Animal communication often relies on various contextual cues to recognize kinship, especially in large groups (Beecher, 1982). Among bird species, oscine songbirds (Passeriformes) are renowned for their vocal learning abilities and have been the primary focus of research on avian communication, contributing significantly to our understanding of neuroanatomical motor control of learned songs (Vicario et al., 2002) and revealing a critical developmental threshold for vocal learning in most oscine songbirds (Tramontin & Brenowitz, 1999). Vocal non-learners, such as seabirds, present a challenge in understanding avian communication because it is believed they rely on innate vocalizations rather than learning their vocalizations through mimicry and practice (Jones & Ryan, 2010). Vocal non-learning has been studied less extensively than vocal learning in birds, and it remains unclear whether vocal non-learners exhibit thresholds for adult vocalizations.

Research on the development of vocalizations in colonial breeding seabirds has primarily focused on how these vocalizations are used in social and environmental contexts (Aubin et al, 2000; Aubin & Jouventin, 2002; Beecher, 1982; Clark et al, 2006; Favaro et al., 2015; Favaro et al., 2016; Jouventin & Aubin, 2002; Kidawa et al, 2023; Lynch & Lynch, 2017; Marks et al, 2010; Robission et al, 1993; Sladen, 1958; Spurr, 1975; Terranova et al, 2023). Such research has shown how vocalizations are essential for mating, territory defense, and offspring recognition (Jouventin & Aubin, 2002). One study suggests that changes in pitch in the vocalizations of Little Blue Penguins (*Eudyptula minor*) were variable between individuals and could act as an identity marker (Nakagawa et al., 2001). However, limited research exists on the vocal development of penguin chicks before they fledge, making it challenging to understand how vocalizations emerge and evolve in the early life stages of colonial seabirds. Here, we aim to test if specific acoustic features in non-passerine chicks change predictably with age as part of developmental growth.

Penguins are considered vocal non-learners, demonstrating limited vocal learning in specific social contexts. For example, breeding adult penguins perform vocal adjustments in pitch, duration, syllables, and rhythm to communicate identity (Robsion et al., 1993; Jouventin & Aubin, 2002). *P. adeliae* perform a loud mutual display call during the breeding season that likely evolved for the purpose of mutual recognition between individuals within loud, congested breeding colonies (Aubin & Jouventin, 2002). Chicks perform a begging peep from the day they hatch, and they have been known to perform immature versions of the loud mutual display call (Davis, 1982; Spurr, 1975). Acoustic modulation in these calls facilitates mate and parent-chick recognition and ensures parental investment is given to the genetic offspring within the colony (Speirs et al., 1991). Mutual vocal recognition is particularly crucial for species with high parental investment and a risk of offspring misidentification (Kidawa et al., 2023). However, the stage at which chicks begin to exhibit vocal modulation patterns or if they participate in vocal learning during brooding remains unclear. Addressing these gaps could improve our understanding of seabird communication and its ecological and evolutionary implications and provide a measure to track potential phenological changes due to climate change in polar regions.

### Chick Age Classes

*P. adeliae* chicks are semi-altricial and require significant parental care. The initial phase, known as the “guard stage,” spans from 0 to 20 days of age (Sladen, 1958). During the first 10 days, chicks are unable to thermoregulate and are immobile, making them highly susceptible to predation (Spurr, 1975). In this stage, both parents alternate between guarding the nest and foraging at sea, with one adult always present to tend to the chicks (Spurr, 1975). On average, the guard stage lasts until the chicks are 20 days old, at which point they become homeothermic, allowing both parents to shift their focus primarily to foraging. Following the guard stage, chicks enter the “crèche stage,” which occurs from approximately 21 to 41 days of age, and is marked by rapid growth (Taylor & Roberts, 1962). During this period, chicks become more mobile and begin to form crèches, aggregations that provide warmth and protection from predators while the parents are at sea (Davis, 1982; Spurr, 1975; Taylor & Roberts, 1962). Selective pressures for individual recognition may be driven by this behavior, enabling parents to identify and retrieve their offspring from the crèche (Kidawa et al., 2023). In this study, the final developmental stage before the chicks fledge is referred to as the “post-crèche” stage. During this stage, chicks exhibit increased independence as they prepare to leave the colony and live at sea. Large groups of chicks gather on beaches to fledge at approximately 54 days post-hatch (Cimino et al, 2014; Salihoglu et al, 2001).

This study explores the vocal development of *P. adeliae* chicks using Passive Acoustic Monitoring (PAM) combined with machine learning. Vocalizations were recorded across different chick developmental stages to examine how these calls change over time. PAM is a minimally invasive, cost-effective data collection method that involves the use of unsupervised, autonomous acoustic sensors to continuously record sounds in challenging or sensitive environments. This method is particularly valuable for long-term monitoring of wildlife distribution and behavior, especially in colonial, highly vocal species where breeding grounds are costly and invasive to access by traditional monitoring methods. However, PAM generates large volumes of data, making manual annotation time-consuming and inefficient. To address the challenge of data volume, machine learning was introduced for efficient processing. Machine learning uses computational models to learn representations of natural sight, vision, and hearing for detection and classification (LeCun et al., 2015). Machine learning, particularly deep learning, offers powerful tools to automate the detection and classification of vocalizations. By leveraging computational models that can learn complex patterns in sound, machine learning significantly reduces the time and effort required to analyze acoustic data, making PAM more efficient and scalable for studying wildlife behavior over extended periods.

## Materials and Methods

Five Adélie penguin colonies of various sizes were continuously recorded during the austral summer from November 2022 – February 2023 near Palmer Station in the West Antarctic Peninsula using Wildlife Acoustics Song Meter Minis (SMMs, Wildlife Acoustics, Maynard, MA, USA). Two colonies were located on Humble Island, and three were located on Torgerson Island. Penguin nests remained in the same location throughout this period, and nests were monitored daily by visual inspection of 30 nests per island.

### Acoustic Recorder Deployment and Programming

Five SMMs were deployed on recording rigs adjacent to *P. adeliae* colonies. Each rig consisted of a PVC structure with a recorder zip-tied at a height of 1 m to keep it out of reach of the penguins and positioned within 1 m of the colony edge (Fig. 1). At Humble Island, two rigs were placed near the colonies due to the size of the colony, while at Torgersen Island, a single rig was positioned near the approximate center of the three colonies, yet still at the edge of the colonies. Rocks were placed at the base of each rig to provide additional stability. The SMMs recorded five-minute audio clips at the start of every hour, sampled at 24 kHz with 16-bit depth and a 6 dB gain. Memory cards and batteries were replaced monthly. Recordings were collected during the *P. adeliae* breeding season, from 27 November 2022 to 27 February 2023, with deployment occurring after the peak egg-laying date to minimize the risk of nest abandonment. In total, 13,932 five-minute recordings were obtained, all wild *P. adeliae* vocalizations. The original audio files were used for all subsequent analyses.

**Figure 1:**
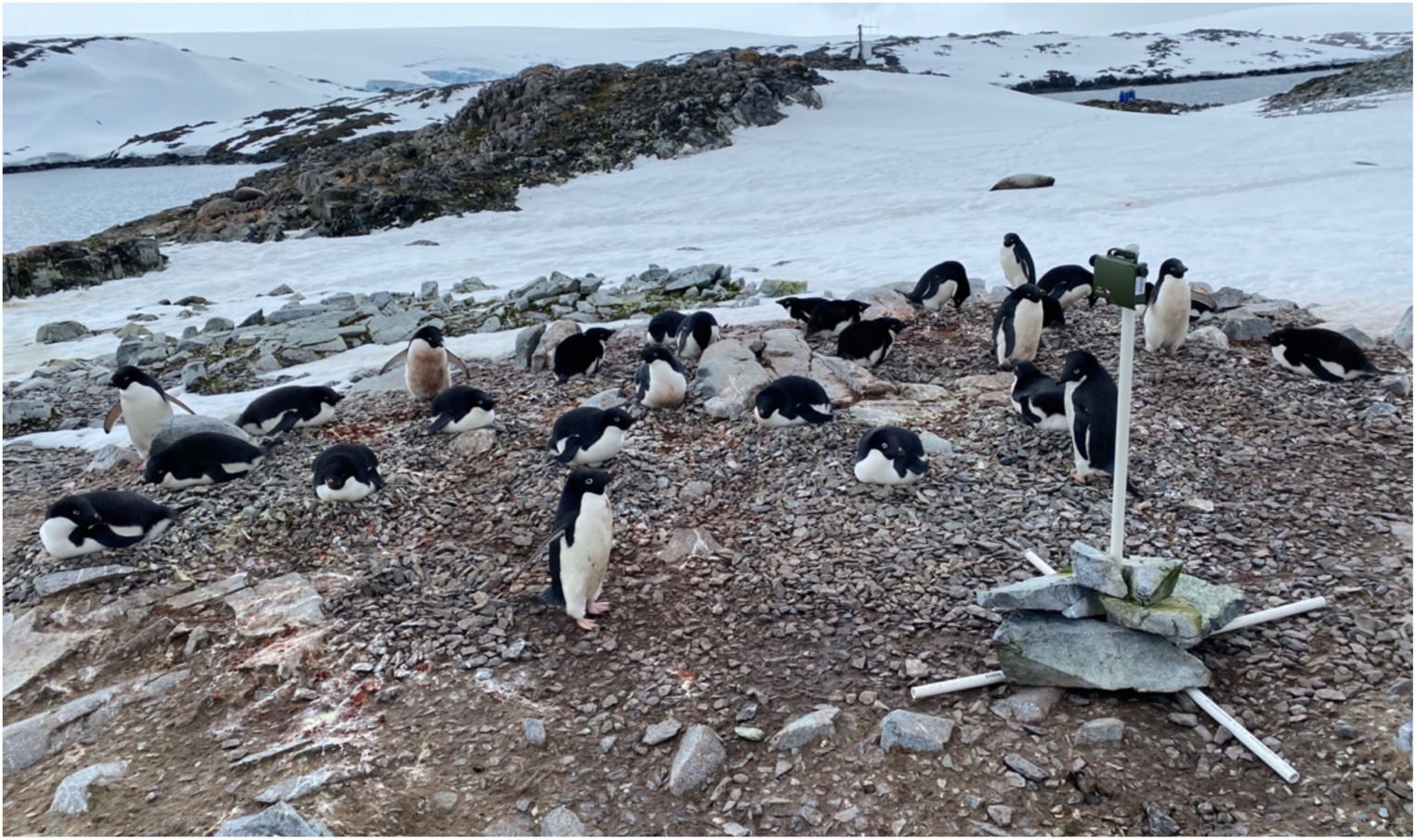
Photograph of recording rig deployed near the edge of an Adélie penguin colony with the acoustic sensor at the top.

### Data Processing

This study employs DeepSqueak, a deep learning-based system, to investigate the vocal development of *P. adeliae* chicks. DeepSqueak is a call recognition software designed for non-expert users to detect, measure, and classify mice and rat vocalizations (Coffey et al., 2019). Because DeepSqueak’s architecture uses spectrogram images, it is important to include different call types into training categories. Prior to training and testing of DeepSqueak, call types were differentiated through manual visual inspection of spectrograms, with particular attention given to identifying chick calls and separating them from adult vocalizations. Calls were broken into three categories based on spectrographic characteristics: begging peep (BEG), multi-syllable beg (MSB), and loud mutual display (LMD). Early spectral BEG calls were identified as very short, high-frequency vocalizations that appeared as small, discrete “carrot-shaped” elements above the frequency range of adult calls. MSB calls were characterized by multi-syllable begging vocalizations that lacked consistent contour structure and showed substantial variability across syllables. In contrast, LMD calls were longer in duration and exhibited a more refined and consistent contour, with repeated syllables forming a stable spectrotemporal pattern. These visually distinct features guided the assignment of calls to training categories, and representative spectrograms of each call type are shown in Fig. 2.

**Figure 2.**
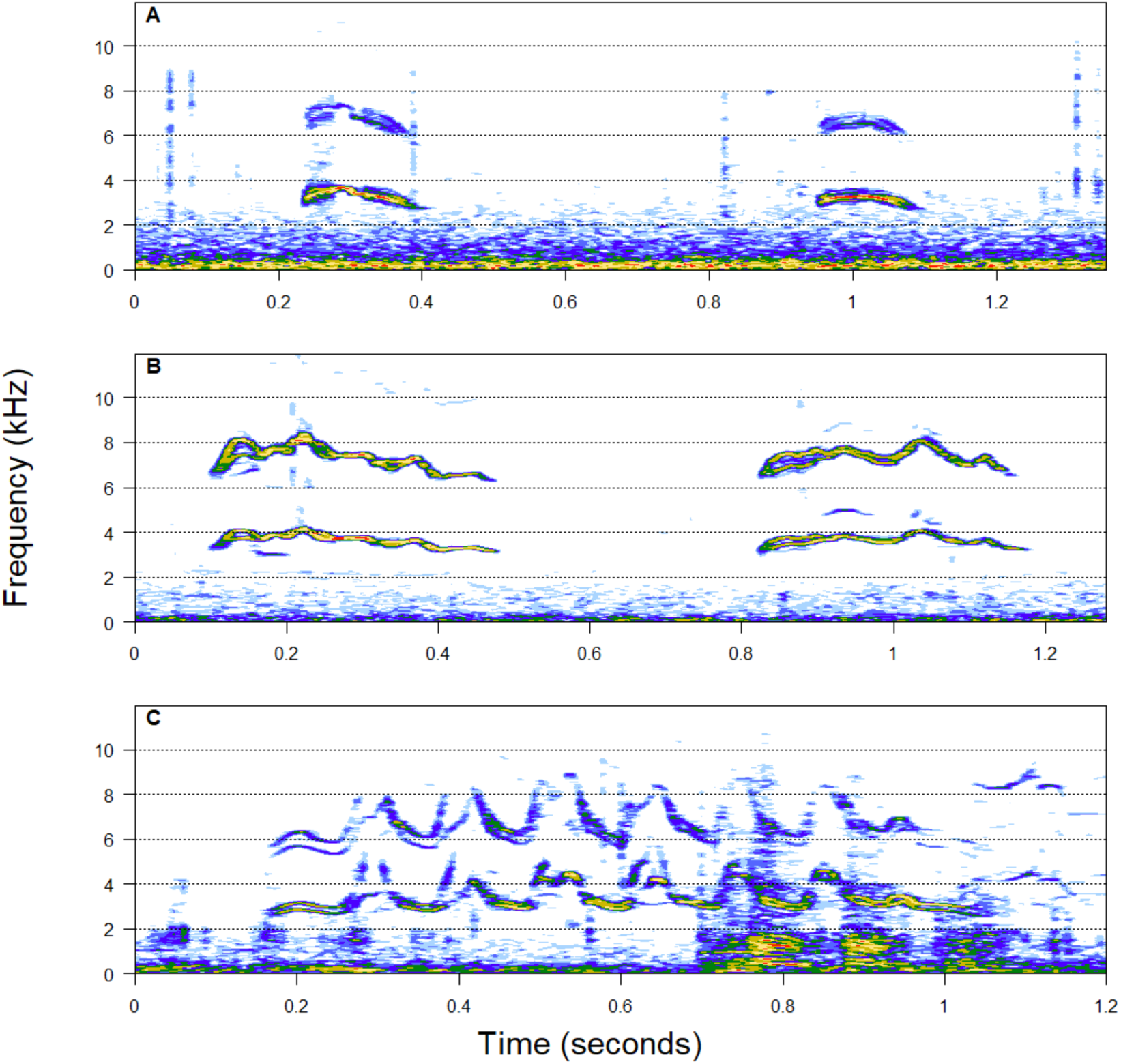
Spectrographic examples of the progression from single syllable begs to multi-syllable *P. adeliae* chick vocalizations highlight the structural changes over times; (a) beg (BEG), (b) multi-syllable beg (MSB) and (c) immature loud mutual display (LMD).

The training dataset consisted of 81 audio files containing *P. adeliae* chick vocalizations. These included 43 BEG recordings, all from the Cornell Macaulay Library; 28 MSB recordings, all obtained from the Macaulay Library; and 10 LMD recordings, of which 2 were sourced from the Macaulay Library and 8 were sourced from our own recordings and excluded from the test data set. Macaulay Library recordings spanned a wide temporal range, with collection dates of 3-5 February 1988, 9 January 1989, 11 January 1989, 23 January 2010, 9 January 2014, 26 January 2022, 8 February 2024, and 5 January 2025. The test data set consisted of 1,848 five-minute segments of our total original recordings, collected between 13 December 2022 and 27 February 2023 and filtered to exclude files where a) chicks were not visually confirmed to be present on the islands and b) recordings where wind noise dominated the soundscape.

After we built our training dataset, we ran our detector on the test data set. Detections were manually reviewed and edited to select the lowest harmonic band for each true positive detection and false positives were labeled within DeepSqueak. For each detection DeepSqueak measured multiple acoustic parameters; the parameters relevant to and reported in our study are listed and described in Table 1. The audio data were divided into three sampling groups distinguished by age class consistent with *P. adeliae* life stage and verified by visual observations: guard stage (1), crèche stage (2), and post-crèche stage (3). The guard stage (weeks 1-3) included all recordings between 13 December 2022 and 02 January 2023. The crèche stage included all recordings between 03 January 2023 and 16 February 2023 (weeks 4-9). The post-crèche stage (week 10) group included all recordings between 17 February 2023 and 27 February 2023, but due to low numbers of chick calls and poor signal-to-noise ratio, we did not include these files in our analysis.

**Table 1.** DeepSqueak acoustic parameters and definitions used in this study.

| Parameter | Definition |
| --- | --- |
| Duration (s) | Total call time, End Time - Begin Time |
| High Frequency (kHz) | Highest frequency of the contour |
| Low Frequency (kHz) | Lowest frequency of the contour |
| Delta Frequency (kHz) | Change in frequency, High Freq - Low Freq; renamed to Bandwidth in our analysis |
| Sinuosity | Length of the path between the first and last points on the contour, divided by the Euclidean distance between the first and last points. |

### Statistical Analyses

To investigate temporal change in chick calls we first applied multinomial regression modeling to quantify developmental shifts in call-type probability across eight weeks. Second, we used a series of generalized additive models (GAMs) to examine how each acoustic feature (bandwidth, duration, high frequency, low frequency, peak frequency, and sinuosity) varied for week, call-type (BEG, MSB, and LMD calls) and the interaction between week and call-type.

To characterize population-level changes in acoustic features of *P. adeliae* chick calls across time, we analyzed all verified detected calls pooled across call types. For each acoustic parameter (bandwidth, duration, high frequency, low frequency, peak frequency, and sinuosity), we fit a generalized additive model (GAM) using week as a continuous predictor. GAMs were implemented using thin-plate regression splines with restricted maximum likelihood estimation to flexibly capture non-linear temporal trajectories while avoiding overfitting. To identify periods of abrupt change in the direction or rate of each trajectory, we applied segmented (piecewise linear) regression to the GAM-predicted values, using week as the segmentation variable. This two-stage approach allowed breakpoint detection to be driven by smooth population-level trends rather than by noise in individual observations. For each acoustic feature, the segmented regression yielded an estimated breakpoint (in weeks) and slopes before and after the breakpoint.

## Results

### Call Type Variation Across Developmental Week

DeepSqueak detected 2579 calls, all of which were manually verified resulting in 1636 true detections (1395 BEG; 111 MSB; 127 LMD). Over the eight weeks of sampling, the composition of call types shifted noticeably throughout the phenological stages (Table 2). BEG calls dominated early in the sampling period (guard stage), comprising nearly all vocalizations in weeks 1 and 2 (>99%), while LMD and MSB calls were initially rare, with proportions below 1%. By week 3 (the transition from guard to crèche stage), the proportion of BEG calls decreased to 58%, while MSB calls increased substantially to 31%, and LMD calls rose to 11%, indicating the emergence of more complex or varied vocalizations. Interestingly, the proportion of BEG calls rebounded in Week 4 to 91%, with LMD and MSB call types remaining low. Later weeks demonstrated further heterogeneity. In week 5, MSB calls (58%) surpassed LMD (42%), while BEG was absent. Weeks 6 and 7 showed BEG calls regaining dominance (71–87%), with LMD and MSB present in smaller proportions. By week 8, BEG calls comprised 70%, and LMD 30%, with MSB absent.

**Table 2.** Proportions of call types from 24 December 2022 to 16 February 2023 shown as the total number of verified detections per call type per week, with the proportion of each call type relative to the total number of calls.

| Week | BEG (n / prop) | LMD (n / prop) | MSB (n / prop) | Total |
| --- | --- | --- | --- | --- |
| 1 | 612 / 0.990 | 2 / 0.003 | 4 / 0.006 | 618 |
| 2 | 290 / 0.990 | 3 / 0.010 | - | 293 |
| 3 | 147 / 0.583 | 28 / 0.111 | 77 / 0.306 | 252 |
| 4 | 48 / 0.906 | 5 / 0.094 | - | 53 |
| 5 | - | 11 / 0.423 | 15 / 0.577 | 26 |
| 6 | 137 / 0.706 | 41 / 0.211 | 16 / 0.083 | 194 |
| 7 | 120 / 0.870 | 17 / 0.123 | 1 / 0.007 | 138 |
| 8 | 41 / 0.695 | 18 / 0.305 | - | 59 |

Overall, BEG calls were consistently the most frequent, but the data indicate a clear temporal trend toward increasing contribution of LMD and MSB calls during the middle of the observation period, followed by a slight return to BEG dominance in the final weeks. These results were reflected in the predicted probabilities of each call from multinomial regression (Fig. 3).

**Figure 3.**
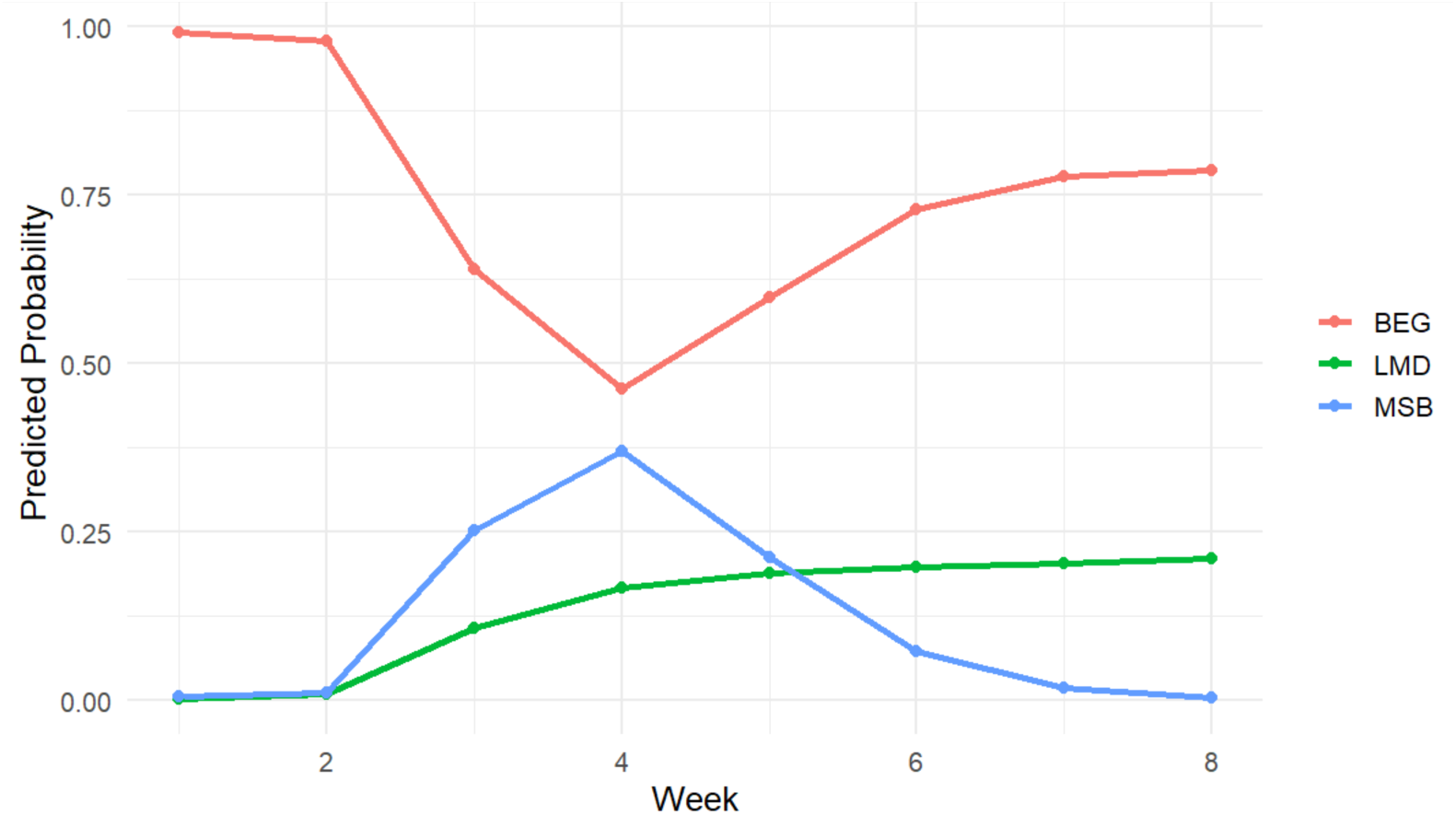
Predicted probabilities of chick call types over the brooding period. Predicted probabilities for each call type (BEG, LMD, MSB) are shown across weeks 1–8, illustrating the changing likelihood of each call type with age. Early development is dominated by BEG calls; mid-development shows greater variability among call types, and later weeks return to higher BEG probability.

### Temporal Trends in Acoustic Features

Changes in acoustic features over time for each call type are shown in Fig. 4. Bandwidth increased over time in BEG (0.268–0.361 kHz) and MSB calls (0.242–0.561 kHz), whereas LMD calls peaked mid-season (0.498 kHz at week 4) before declining to 0.337 kHz by week 8. Duration showed strong temporal increases in BEG (0.153–0.302 s) and MSB calls (0.220–0.552 s), while LMD calls exhibited higher overall durations early in development (0.802 s at week 1) followed by a gradual decline toward week 8 (0.597 s). Consistent with high model fit (Table 3), duration showed the strongest temporal structure across call types.

**Figure 4.**
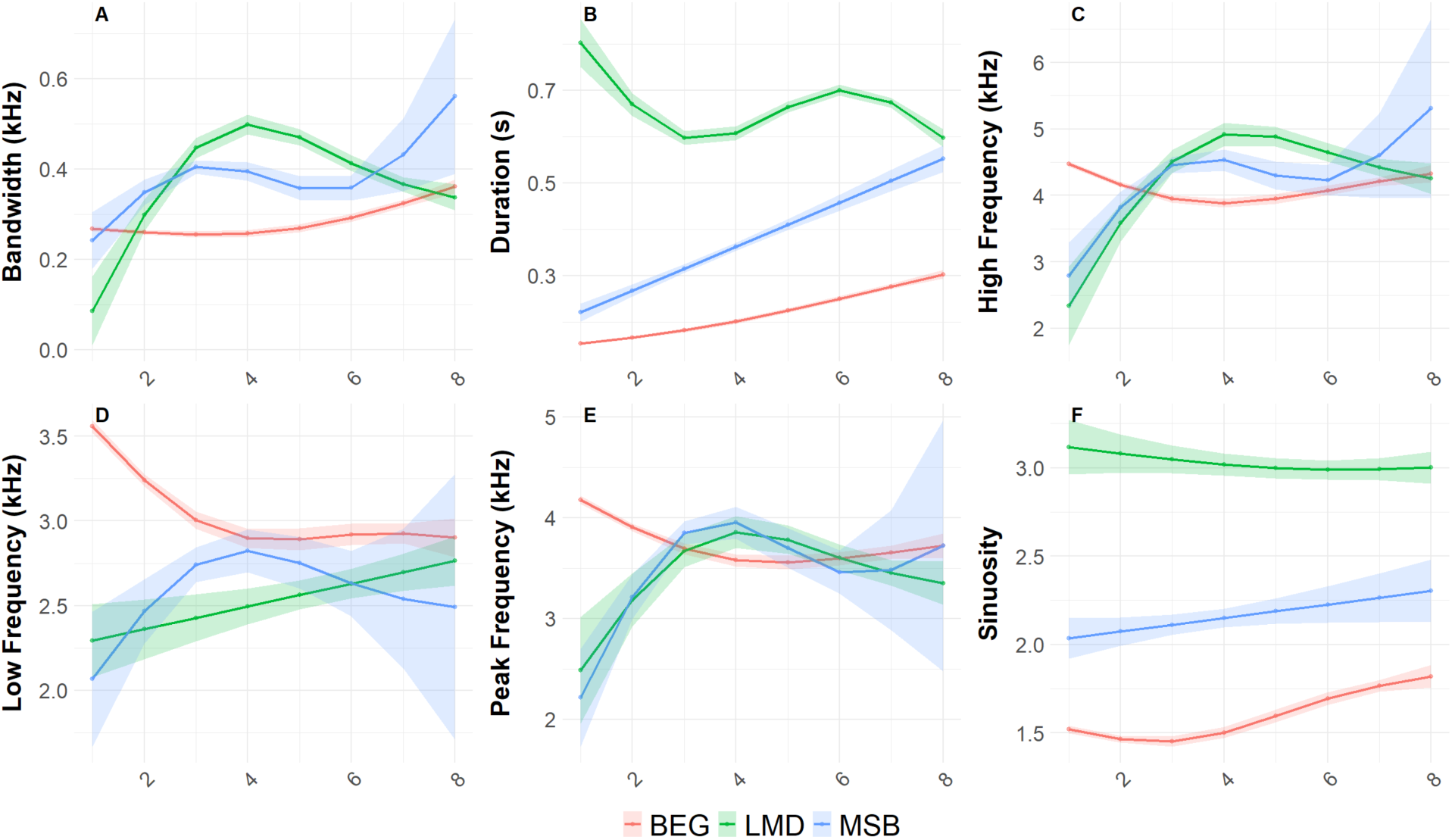
Developmental trajectories of acoustic features in chick calls. GAM-derived fitted values (± SE) are shown across weeks 1–8 on the x-axis for three call types (BEG, LMD, MSB) for each acoustic feature: bandwidth, duration, high frequency, low frequency, peak frequency, and sinuosity.

**Table 3.** Model fit statistics for generalized additive models (GAMs) describing temporal variation in acoustic features across chick development. Temporal patterns indicate whether nonlinear change across weeks was detected consistently across call types or restricted to specific call types.

| Acoustic feature | n | Adjusted $R^2$ | Deviance explained (%) | Temporal Pattern Detected |
| --- | --- | --- | --- | --- |
| **Bandwidth** | 1633 | 0.135 | 14.1 | All call types |
| **Duration** | 1633 | 0.705 | 70.7 | All call types |
| **High frequency** | 1633 | 0.046 | 5.23 | All call types |
| **Low frequency** | 1633 | 0.117 | 12.1 | BEG only |
| **Peak frequency** | 1633 | 0.062 | 6.77 | BEG, MSB |
| **Sinuosity** | 1633 | 0.384 | 38.6 | BEG only |

All models used a Gaussian error distribution with identity link. Temporal variation was modelled using penalized smooths of week fitted separately for each call type ( $s(\text{week}, \text{by} = \text{call\_type}), k = 4$ ). “Temporal pattern detected” summarizes whether the smooth term of week was significant ( $p < 0.05$ ) for all call types or only a subset, reflecting call-type-specific temporal structure rather than overall differences among call types.

High-frequency components were relatively stable in BEG calls (3.88–4.33 kHz), increased substantially in MSB calls (2.79–5.31 kHz), and peaked mid-season in LMD calls (4.92 kHz at week 4) before declining. Low-frequency components decreased slightly in BEG calls (3.56–2.90 kHz), increased gradually in LMD calls (2.29–2.76 kHz), and showed a mid-season peak in MSB calls. Although these frequency-related features exhibited weaker overall model fit, significant temporal effects were detected for selected call types (Table 3). Peak frequency remained largely stable in BEG calls, while MSB and LMD calls exhibited mid-season peaks. Sinuosity increased gradually over time in BEG calls (1.52–1.82) and MSB calls (2.03–2.30), whereas LMD calls maintained consistently higher sinuosity values with little temporal change. Consistent with model-fit results, significant nonlinear temporal change in sinuosity was detected only for BEG calls, possibly indicating seasonal modulation of call structure rather than differences in overall complexity.

### Breakpoint Analysis (Segmented Regression)

Recognizing that the transition between call types may be gradual, especially the transition between BEG and MSB, we pooled all call categories together and used a breakpoint analysis to identify major shifts in the vocal behavior during chick development. Segmented regression identified significant shifts in the rate of change for each acoustic feature (Fig. 5). The greatest change in frequency parameters (high frequency, low frequency, bandwidth) occurred during weeks 3 and 4 (crèche stage), whereas the greatest change in duration and sinuosity occurred around week 5 (also in crèche stage).

**Figure 5.**
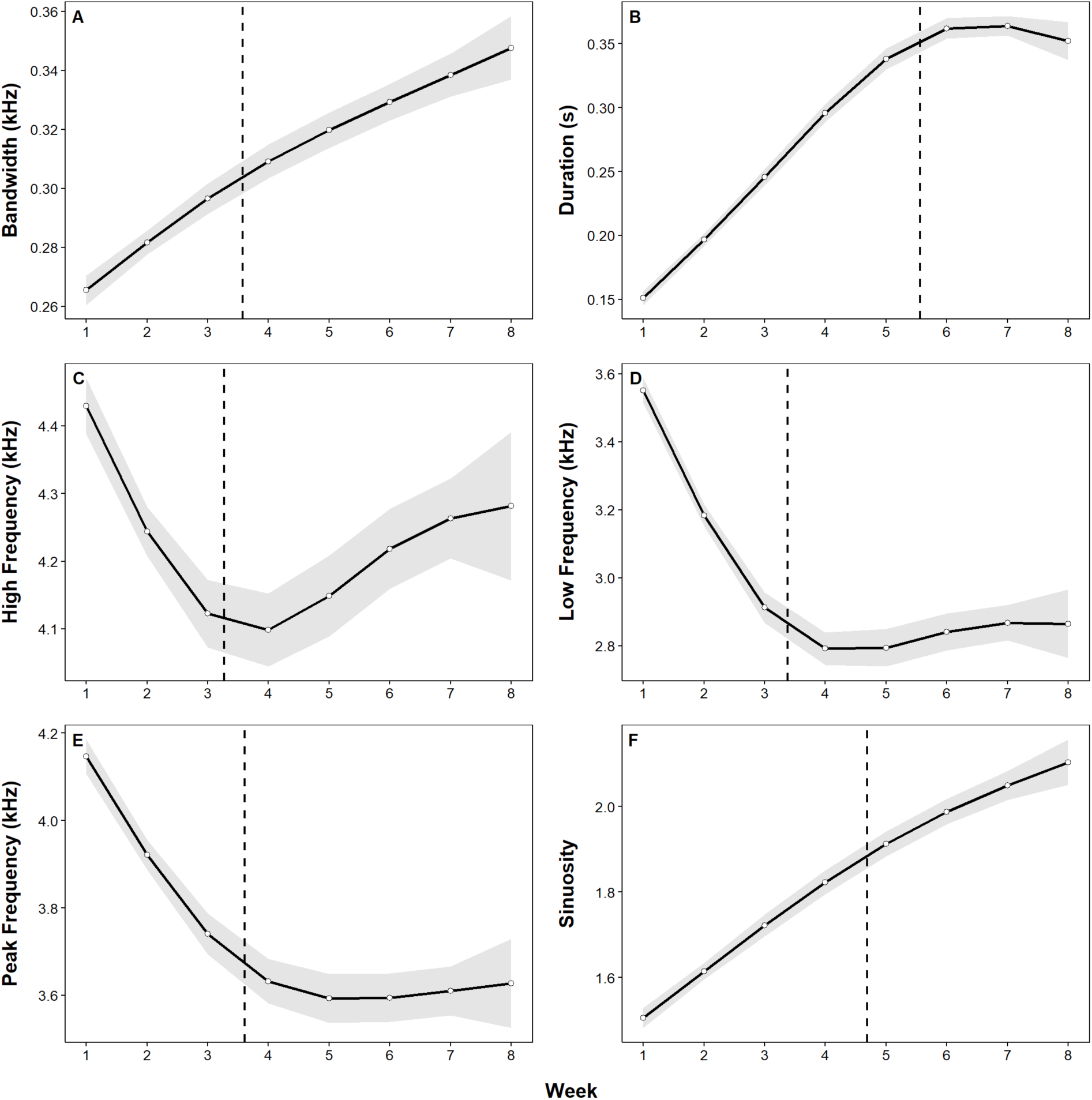
Breakpoint analysis of chick call features. Data represents all call types pooled together. Vertical lines indicate weeks where segmented regression identified a significant shift in the rate of change for each acoustic feature, highlighting potential critical periods of vocal development. Shaded regions illustrate the slope before and after each breakpoint, with corresponding values provided in Table 4.

**Table 4.** Breakpoints were estimated using segmented linear regressions fitted to GAM-predicted values averaged across call types. Slopes represent the rate of change per week before and after the estimated breakpoint.

| Feature | BP Week | Slope before BP | Slope after BP | Directional change |
| --- | --- | --- | --- | --- |
| **Bandwidth** | 3.57 | 0.016 | 0.01 | Increase → slower increase |
| **Duration** | 5.56 | 0.047 | −0.005 | Increase → plateau / slight decline |
| **High freq.** | 3.27 | −0.154 | 0.048 | Decrease → increase |
| **Low freq.** | 3.38 | −0.320 | 0.022 | Decrease → increase |
| **Peak freq.** | 3.61 | −0.203 | 0.001 | Decrease → stabilization |
| **Sinuosity** | 4.69 | 0.106 | 0.064 | Increase → slower increase |

## Discussion

As *P. adeliae* chicks develop, their vocalizations also change in both temporal pattern and acoustic features. During the guard stage, vocalizations are overwhelmingly BEG calls, characterized by short (< 0.3 seconds) durations, higher (4.5 kHz) frequencies, and low sinuosity. As the chicks transition to the crèche stage, MSB and LMD calls begin to emerge, characterized by longer durations, lower frequencies, and more sinuosity compared to the BEG calls (Fig. 2). MSB predicted probabilities peak around week 4 (start of crèche) with an associated decline in BEG call probabilities, whereas LMD predicted probabilities steadily increase over development (Fig. 3). Within BEG and MSB call types, we see further structural changes as chicks age. Across our observed development period, BEG and MSB calls became more broadband, longer, and more tonally complex (sinuosity), while LMD calls peaked in bandwidth around week 4 and demonstrated consistent sinuosity across weeks (Fig. 4). Together, these patterns suggest progressive modification of call structure across developmental stages, with some call types showing more pronounced temporal changes than others. These changes may be related to colony dynamics due to developmental changes, behavioral transitions, and/or shifts in colony activity that favor specific call types. For example, during the guard stage, when the chicks are primarily near their parents, BEG calls would be expected, as these calls are used to stimulate the parent to feed the chick (Spurr, 1975). In contrast, during the crèche stage, chicks are unguarded by parents for several hours a day and left in clusters with other chicks. Upon reunion with parents, both “immature versions of the adult LMD” and complete LMD are made prior to begging calls, which play a crucial role in parent-offspring recognition (Spurr, 1975), with this recognition emerging between 8 and 17-days post-hatch (Thompson and Emlen, 1968). Our results are consistent with the description from Spurr (1975), especially if our classified MSB is an immature version of the LMD. The similarities in spectral features (Fig. 2) and the peak in MSB at the start of crèche (Fig. 3), suggest the MSB should be considered a precursor to the LMD. The MSB/LMD reunification calls occur with a behavioral display clearly distinguishable from that of begging (Spurr 1975).

Breakpoint analysis further reinforces the suggestion that changes in the chick call features are associated with developmental stage. All spectral features (bandwidth, high frequency, peak frequency, low frequency), demonstrate a significant shift in the rate of change between weeks 3 and 4, which is at the start of the crèche period. This is consistent with the emergence (Fig. 3) and subsequent stabilization of frequency for the MSB and LMD calls (Fig. 4), suggesting early refinement of critical call spectral features, which may be important for parental recognition. Although the spectral features of calls remain relatively stable after week 4, the duration and sinuosity of calls continue to increase over development, reflecting later enhancement of complex calls possibly used for social signaling or competition, which may be significant during the crèche stage.

In “open-ended” vocal learners such as songbirds (passerines), high vocal sinuosity in early developmental stages have represented exploratory or “babbling” behavior as a way for juveniles to practice sounds before vocal motor control is refined (Arato et al, 2021; Elie & Theunissen et al, 2020; Aronov et al., 2008). As they mature and experience vocal plasticity, the variability in sinuosity decreases and aligns with increased control and stability of sound (Arato et al, 2021). This process is characteristic of species that undergo a distinct learning phase via practice, feedback mechanisms, and maturation of vocal organs (Elie & Theunissen et al, 2020). However, in “closed-ended-learners” (i.e., species with limited vocal modification potential) that rely on innate vocal patterns, vocal development may not rely on such high variability (Arato et al, 2021). While sinuosity cannot capture the combined effect of phenological stage and call type together, it may provide valuable ontological information when the context of each factor is interpreted separately. In this context, an increase in sinuosity with age could suggest an increase in control over vocalizations, suggesting vocal refinement in older chicks. Conversely, very low sinuosity in younger chicks may reflect less control or simple vocal structures. In contrast to the vocal development of passerine birds, these data may suggest that high sinuosity corresponds with more complex or modulated calls in the penguin chicks, as opposed to indicating refined, less variable calls in passerine species. In other words, the complexing of learned avian *song* may decrease to signify mature vocalizations, whereas the complexing of avian *calls* in non-passerines may increase to signify mature vocalizations.

For our analysis, we grouped calls based on call types from distinct spectrographic features, but there is the possibility that these call types are instead the same call in different forms. When we accounted for this by grouping all call types together, we still see shifts in vocal parameter across weeks (Fig. 5), indicating chicks undergo gradual vocal development as they age. This could suggest developmental fine-tuning within a fixed call type, supporting a form of vocal maturity without the acquisition of new calls. If the call functions primarily for a limited purpose, like signaling hunger or distress, they will not require complex vocal repertoires at early developmental ages. However, we know that *P. adeliae* adults modulate calls in courtship rituals and identification (Aubin & Jouventin et al, 2002; Clark et al, 2006) and descriptive studies of chicks indicate multiple call types associated with different behaviors (Spurr, 1975). We therefore propose that even in early developmental stages, chicks use different call types, with the emergence of modulated calls (MSB and LMD) occurring at the start of the crèche.

Beyond our study period, there is likely further refinement of call modulation as chicks transition their LMD calls to their adult form. In adult *P. adeliae,* LMDs are individually distinctive and play a key role in identifying mates or communicate to conspecifics. Males discriminate more than females to identify individuals and are more territorial with higher fidelity to offspring than females (Speirs et al., 1991). Given the role the LMD plays in individual recognition, learning likely plays a role, supporting the hypothesis that simpler vocal learning mechanisms may exist outside the current understanding of vocal learning in birds (ten Cate, 2021; Baciadonna et al., 2022) and underscoring the need for more diverse models than those used for the complex vocal development seen in passerine birds (Vernes et al., 2021). Given the role that physical displays play with the associated vocalizations (Spurr, 1975), further analysis could incorporate behavioral elements to investigate whether specific social contexts lead to different call patterns, potentially indicating some degree of plasticity. For instance, does the presence or absence of parents or other adults near the nest affect the call rate or acoustic features of chick vocalizations? Additionally, what role does sinuosity play in adult courtship vocalizations, and how might sinuosity and call duration relate to chick survival or reproductive success in subsequent years?

### Sources of Error and DeepSqueak Utility

Our analysis also relies on general knowledge of breeding phenology and timelines supported by literature (Cimino et al., 2023; Spurr, 1975; Taylor & Robertson, 1962) and visual observations, but it is important to note that these timelines are approximate, and visual observations can be imperfect, especially considering not every nest is monitored. Additionally, we categorized our calls based on unique spectrographic differences, but some call types may be inherently more variable or transitional, such as the MSB call, making them more prone to detection errors. This could introduce biases in observed patterns, both in DeepSqueak’s detection networks and under variable environmental conditions, such as wind. For example, wind may artificially broaden measured bandwidth, potentially confounding true biological changes with recording artifacts, or affect penguin vocal behavior (Zhao et al., 2022). The 2022-2023 Antarctic summer was an extremely windy period (Fradet et al., 2025), so continued recording of chick vocalizations with larger sample sizes can further test the patterns we observed with our data.

The performance of DeepSqueak in detecting chick calls depends largely on user experience and on the supplemental programs used alongside it. Even so, its object-detection framework makes it an appealing option for non-experts, especially compared to more specialized, expert-driven tools that lack this feature. For users seeking both call detection and extraction of acoustic variables, DeepSqueak provides a straightforward and cost-effective solution. Nevertheless, our detection networks encountered some challenges. A single network could not reliably distinguish adults, chicks, and other species, nor could it separate detections by age class when trained on sampling dates rather than observed spectrographic differences among chick calls. Our networks focused on specific call types, identified through contour characteristics in spectrograms, and performed more consistently for detecting *P. adeliae* chick calls. The unsupervised classifier also struggled to group chick calls, likely due to minimal variation among a large number of similar data points.

## Conclusion

These findings demonstrate the development of vocal features during early *P. adeliae* development. In the first few weeks, chicks produce primarily begging calls, characterized by short-duration, low sinuosity, and high frequencies. Around the start of the crèche period, chicks begin to produce multi-syllable beg calls, which may be a precursor to the loud mutual display calls. Over development, acoustic features of calls continue to change, suggesting vocal refinement and a shift to producing signals longer in duration and with more frequency modulation compared to early development. These results raise the possibility of a vocal learning mechanism in penguins, but additional data spanning other breeding seasons and penguin species is needed to continue this investigation.

## Acknowledgements

This project is funded by the National Science Foundation EAGER Award 2226886 and National Science Foundation OPP 2026045. The authors would like to thank Darren Roberts and Megan Roberts for collecting acoustic data and conducting colony surveys, and Kevin R. Coffey for guidance with DeepSqueak. Work was approved by the Antarctic Conservation Act permit #ACA 2021-002.

